# Quantitative analysis of the splice variants expressed by the major hepatitis B virus genotypes

**DOI:** 10.1101/2020.08.12.249060

**Authors:** Chun Shen Lim, Vitina Sozzi, Peter A. Revill, Chris M. Brown

**Author notes:** Equal contributions (Peter A. Revill, Chris M. Brown). Address correspondence to Chris M. Brown, or Peter A. Revill.

## Abstract

Hepatitis B virus (HBV) is a major human pathogen that causes liver diseases. The main HBV RNAs are unspliced transcripts that encode the key viral proteins. Recent studies show that some of the HBV spliced transcript isoforms are predictive of liver cancer, yet the roles of these spliced transcripts remain elusive. Furthermore, a total of 9 major HBV genotypes were isolated from discrete geographical regions of the world, it is likely that these genotypes may express a broad variety of spliced transcript isoforms. To systematically study the HBV splice variants, we transfected the human hepatoma cells Huh7 with 4 HBV genotypes (A2, B2, C2, and D3), followed by deep RNA-sequencing. We found that 12-25% of HBV RNAs were splice variants, which were reproducibly detected across independent biological replicates. This accounted for a total of 6 novel and 6 previously identified splice variants. In particular, 2 highly abundant novel splice variants, in which we called the putative splice variants 1 and 5 (pSP1 and pSP5), were specifically expressed at high levels in genotypes D3 and B2, respectively. In general, the HBV splicing profiles varied across the genotypes except for the known spliced pgRNAs SP1 and SP9, which were present in all 4 major genotypes. Counterintuitively, these singly spliced SP1 and SP9 had a suboptimal 5′ splice site, suggesting that splicing of HBV RNAs is tightly controlled by the viral post-transcriptional regulatory RNA element.

**IMPORTANCE:** HBV infection affects over 257 million people worldwide. HBV is a major cause of liver diseases including cancer and there is no cure. Some HBV RNAs are spliced variants and their roles are largely unclear, although some splice variants have been previously found to be associated with liver cancer. HBV exists as 9 genotypes worldwide with marked differences in replicative capacity and disease sequelae. Whether HBV splice variants vary for the different genotypes is yet to be investigated in depth. Here we sequenced RNAs from 4 major HBV genotypes using a cell culture system. We found 6 new and 6 previously known splice variants across these genotypes. Some novel splice variants were present at high levels, suggesting they could be functionally important. Interestingly, although HBV has adapted to human hosts for over 50,000 years, the most frequently spliced location shared little flanking sequence similarity with that of humans.

## INTRODUCTION

Hepatitis B virus is a common human pathogen that is a major cause of liver cirrhosis and liver cancer. The genomic DNA of HBV is approximately 3.2kb, but can be transcribed into the greater than genome length pregenomic RNA (pgRNA) and preC RNA (pcRNA) (1). The pgRNA encodes the core (C) and polymerase (P) proteins, whereas the pcRNA encodes the pre-core (PC) protein that is subsequently processed into the Hepatitis B e Antigen (HBeAg). The HBV genome is also transcribed into several subgenomic transcripts, namely the preS1, preS2, S, and X mRNAs. The preS1, S2, and S mRNAs encode the 3 surface (S) structural proteins of the HBV particles and subviral particles (HBsAg). The X mRNA encodes the HBx protein.

Many strains of HBV have arisen from distinct geographical distributions of the world. This is partly due to the long history of virus-host coevolution (over 50,000 years) and the lack of proofreading function of the viral reverse transcriptase (2–4). These strains were grouped into 9 major genotypes (A to I) and putative J, and about 30 sub-genotypes (4, 5). There are marked differences in replication phenotype and disease natural history across HBV genotypes (6, 7), yet the pathogenicity of different HBV genotypes and their implications for treatment are still not fully understood. For example, it is possible that severe liver injury caused by genotype C is related to its high replication capacity (8), and/or its more frequent mutations at the basal core promoter and pre-core regions (9, 10).

In addition, different HBV genotypes may produce distinct spliced transcript isoforms whose precise roles are largely unknown (11–14). At least 17 spliced transcripts of pgRNA (14–26) and 4 spliced transcripts of preS2/S (27, 28) were identified in various sources including liver, serum, and transfected cells. Interestingly, a recent study showed that HBV RNA splicing is more efficient in human hepatoma cells than other tested cell-types (14). Furthermore, spliced pgRNA SP1 is the most commonly detected (16, 17, 19, 21, 29–33), although SP3 and SP9 have also been commonly observed in some studies (13, 14, 26).

Notably, HBV splice variants can be encapsidated to form defective viral particles, with replication and envelopment requiring polymerase and envelope proteins supplied *in trans* by wild-type HBV (18, 20, 21, 34). The SP1 transcript also encodes the Hepatitis B Spliced Protein (HBSP) (29), as well as a truncated (by 1 amino acid) PC p22 protein that inhibited wild-type HBV replication by interfering with wild-type capsid assembly (35). The HBSP is a fusion product of the first 46 amino acid residues of the P protein and 47 amino acid residues from a distinct reading frame. A recent breakthrough study showed that HBSP could reduce liver inflammation *in vivo* (33). Three other splice variants that have coding potential are SP7, SP10 and SP13. SP7 encodes the Hepatitis B Doubly Spliced Protein (HBDSP), a putative pleiotropic activator (36), whereas SP13 encodes the Polymerase-Surface Fusion Protein (P-S FP), a structural protein that could substitute the large HBV surface protein (37). This fusion protein could inhibit HBV replication and may play a role in persistent infection. Interestingly, SP10 could also act as a functional RNA that reduces wild-type HBV replication through interaction with TATA box binding protein (38).

An increasing number of studies has shown that the HBV splice variants are associated with the development and recurrence of hepatocellular carcinoma (32, 39, 40), and poor response to interferon treatment (24). Therefore, we aimed to utilise RNA-seq on cells that had been transfected with replication competent clones of different HBV genotypes to (i) quantify the composition of splice variants at the RNA level, (ii) investigate the effects of sequence variations on splicing efficiency, (iii) determine the usage of splice sites, and (iv) understand the host response to viral replication across the major HBV genotypes A to D.

## RESULTS AND DISCUSSION

### Six of 12 HBV splice variants detected were novel transcripts

HBV genotypes A to D expressed a large proportion of spliced transcript isoforms, representing 12-25% of the HBV transcriptomes detected (Fig 1A), showing that HBV splicing was indeed widespread across the genotypes. HBV genotype B2 expressed the highest level of HBV transcriptome, followed by A2, C2, and D3 [5830, 5365, 4783, and 4286 TPM (Transcripts Per kilobase Million mapped reads), respectively; see also Supplementary Fig S1 and Table S1 in read counts].

**Fig 1.**
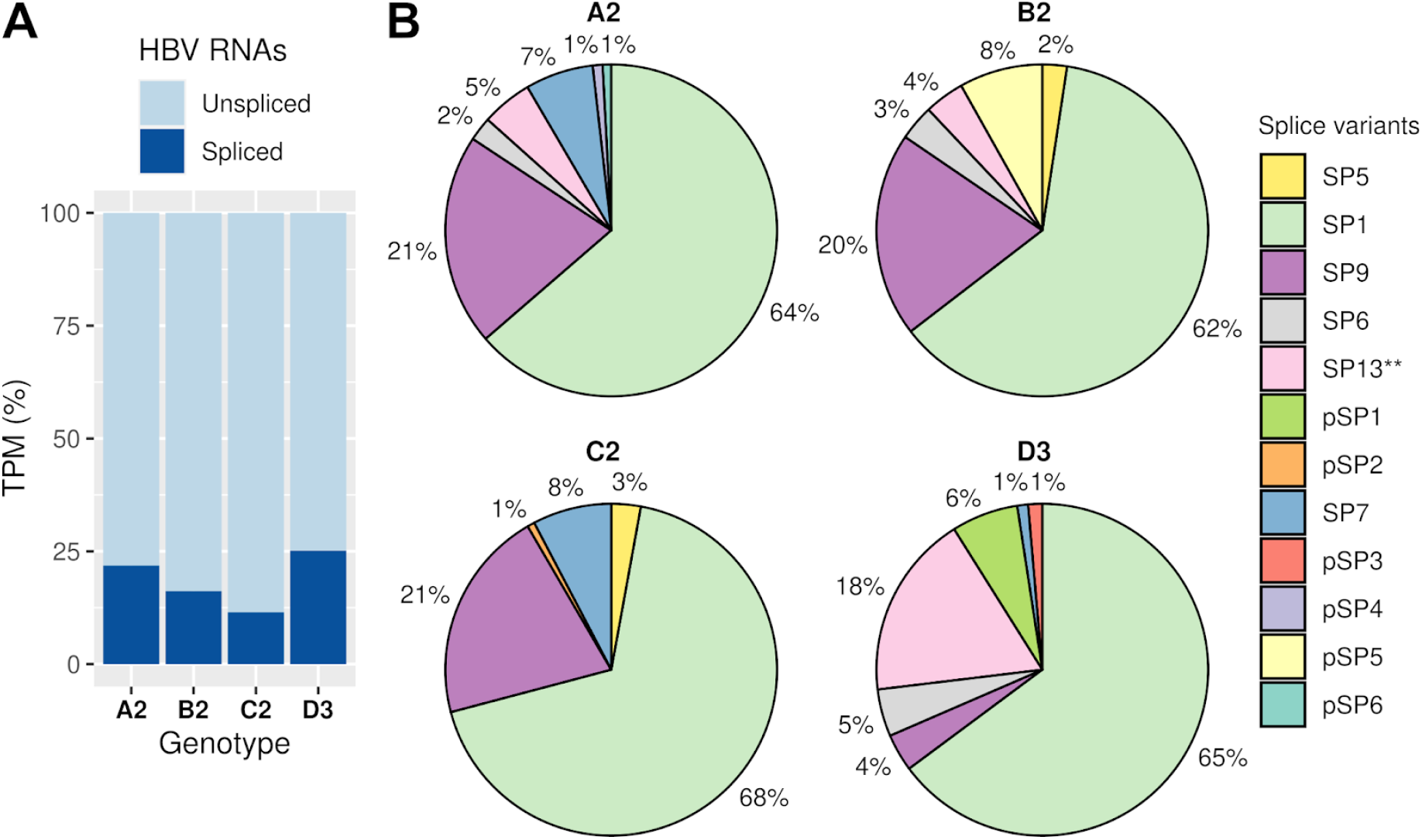
HBV genotypes expressed a wide variety of spliced transcript isoforms. **(A)** Proportions of the HBV spliced transcripts. Only the spliced transcripts present in both biological replicates are shown. **(B)** Relative abundance of the HBV splice variants in genotype A to D. See also Supplementary Table S1 and S2. TPM, Transcripts Per kilobase Million mapped reads.

A total of 12 splice variants were consistently detected across independent biological replicates with high confidence (Fig 1B). In particular, we only reported the splice variants with complete, exact match intron chains across independent biological replicates, which were supported by over 10 reads mapped across the splice junctions, indicating spliced reads (Fig 2, and Supplementary Table S2 and S3). Six out of 12 splice variants were novel. In particular, 2 putative splice variants, pSP1 and pSP5 were expressed at high levels in genotypes D3 and B2, respectively. The importance of these 2 highly abundant novel splice variants requires further investigation.

**Fig 2.**
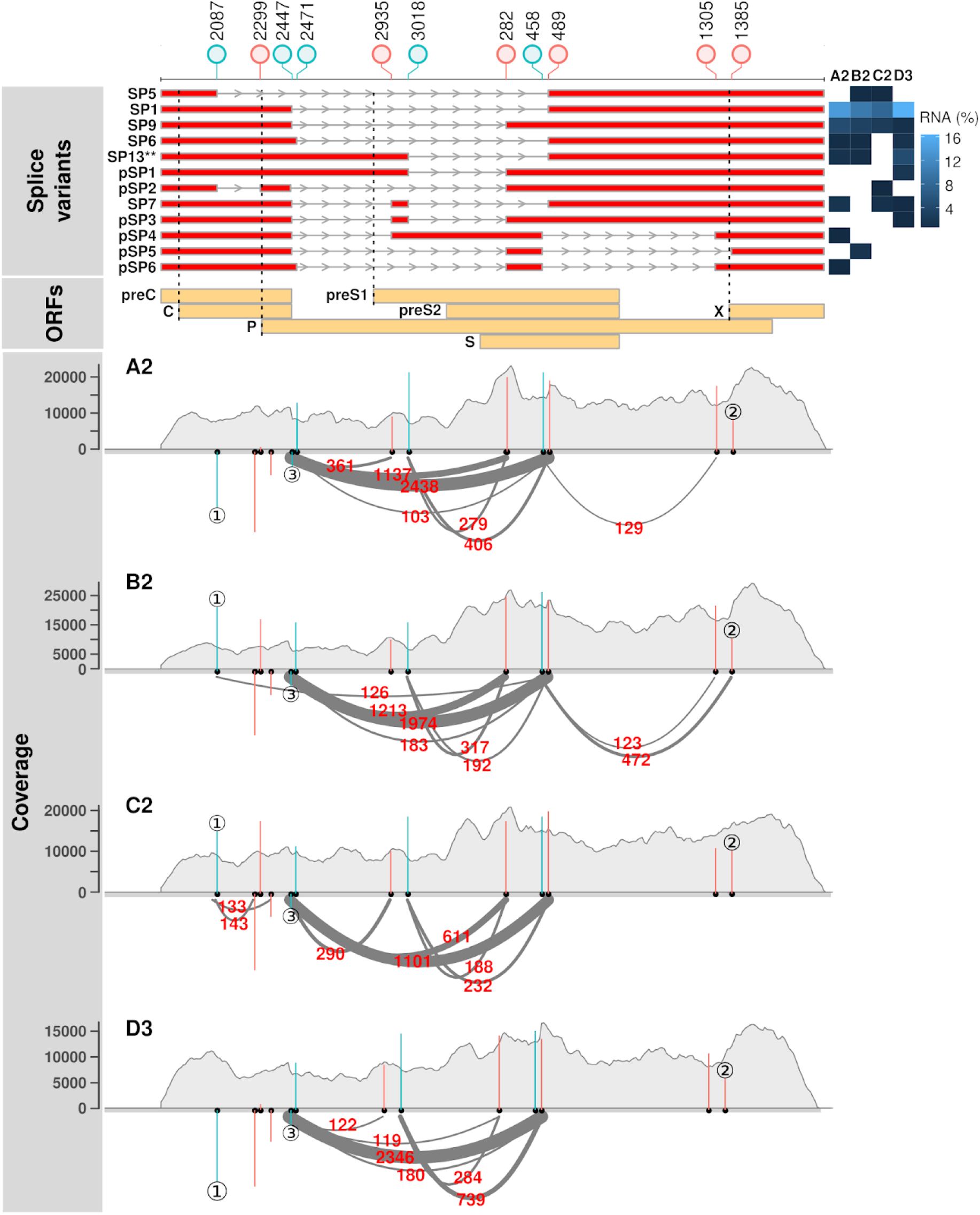
Distinct splicing profiles were observed across the HBV genotypes. Lollipop plot indicates the positions of splice sites relative to the *EcoR*I site of genotype C2. Blue and red colors indicate 5′ and 3′ splice sites, respectively). SP and pSP denote the known and putative spliced pgRNA transcripts, respectively (splice variants’ panel). Black dotted lines denote the positions of initiation codons of C, P, preS1, and X reading frames (ORFs’ panel). Heatmap shows the abundance of HBV spliced transcripts. Read coverage is shown in gray (coverage panel, y-axis). Arcs represent RNA-seq reads mapped across the splice junctions (supporting read counts in red color). Only the splice junctions supported by at least 100 reads are shown. Blue and red vertical lines indicate the MaxEntScan scores of the 5′ and 3′ splice sites, respectively (coverage panel). A positive MaxEntScan score predicts a good splice site sequence context whereas a negative score predicts a poor splice site sequence context. Three main scenarios were observed. ① The presence and absence of spliced reads at position 2087 were predicted by MaxEntScan scores, in which reads were found to map across the 5′ splice sites with strong positive scores (B2 and C2) but not those with strong negative scores (A2 and D3). ② Varying spliced read counts could not be explained by similar scores. ③ Most spliced reads were mapped across a weak splice donor site. See also Supplementary Table S3. ORFs, open reading frames.

We also identified previously reported splice variants SP1, 5, 6, 7, 9 and 13** across the HBV genotypes (11, 14, 24, 38) (Fig 1B and 2). Notably, SP1 and SP9 were consistently detected in all the genotypes. SP1 was the major spliced transcript detected, ranging from 7 to 16% of the HBV transcriptomes, which is in agreement with previous findings (11, 13, 41). SP9 was the second most abundant spliced transcript, ranging from 1 to 5% of the HBV transcriptomes. However, the role of SP9 is still unclear. SP13** was previously identified as a novel splice variant, and we were able to detect it across genotypes A2, B2 and D3 at high levels. In addition, SP6 and SP7 were detected in 3 major genotypes (A2, B2 and D3, and A2, C2 and D3, respectively). While any function is yet to be attributed to SP6 or SP13**, SP7 encodes a putative pleiotropic activator (HBDSP) that has been shown to increase replication of wild-type HBV in co-transfection cell culture experiments (36).

### Sequence variations surrounding the HBV splice sites affected splicing efficiency

We next investigated whether the sequence contexts of splice sites contributed to the different types and abundance of splice variants observed across genotypes. MaxEntScan scoring of the HBV splice sites showed that sequence variation affected the strength of the splice sites of the different HBV genotypes (Fig 2 and Supplementary Table S3).

In general, splice sites with weak sequence contexts (negative MaxEntScan scores) were less likely to be used for splicing and vice versa. For example, the splice donor site at position 2087 in genotypes A2 and D3 had poor sequence contexts and spliced reads associated with this site were not detected (see ① in Fig 2 and Supplementary Table S3). In contrast, the same donor position in genotypes B2 and C3 had strong sequence contexts and were supported by over 100 spliced reads. This indicates that the splicing efficiencies of the HBV RNAs are strongly influenced by the HBV sequence variants surrounding the splice sites.

However, we also observed a discordance between splice site sequence contexts and splice read counts. For example, all the genotypes had similar scores at the splice acceptor position 1385 but the splice read counts were markedly different (see ② in Fig 2 and Supplementary Table S3). In particular, the most frequently used 5′ splice site which was used for SP1 and SP9 had a negative MaxEntScan score (see ③ or position 2447 in Fig 2 and Supplementary Table S3). These results suggest that the splicing of this splice junction may be controlled by other *cis*-regulatory elements, such as the HBV post-transcriptional regulatory RNA element (PRE) (1). Indeed, deleting a PRE component called the splicing regulatory element-1 (SRE-1) was previously found to inhibit pgRNA splicing and the production of SP1 (41). Regulation of alternative splicing may play a crucial role in viral-host interactions (42).

### HBV encoded more alternative 3′ splice sites than 5′ splice sites

A closer examination of the HBV splicing profiles revealed that HBV encoded more alternative 3′ splice sites than 5′ splice sites. (Fig 3A and Supplementary Table S3). Indeed, a trend was observed for more spliced reads mapped across the 5′ splice sites than 3′ splice sites, which reached statistical significance for HBV genotype B2. In contrast, host RNAs had balanced numbers of 5′ and 3′ splice sites (53,009 and 52,998, respectively), as well as the supported read counts (Fig 3, median read counts of 87 for both the 5′ and 3’ splice sites).

**Fig 3.**
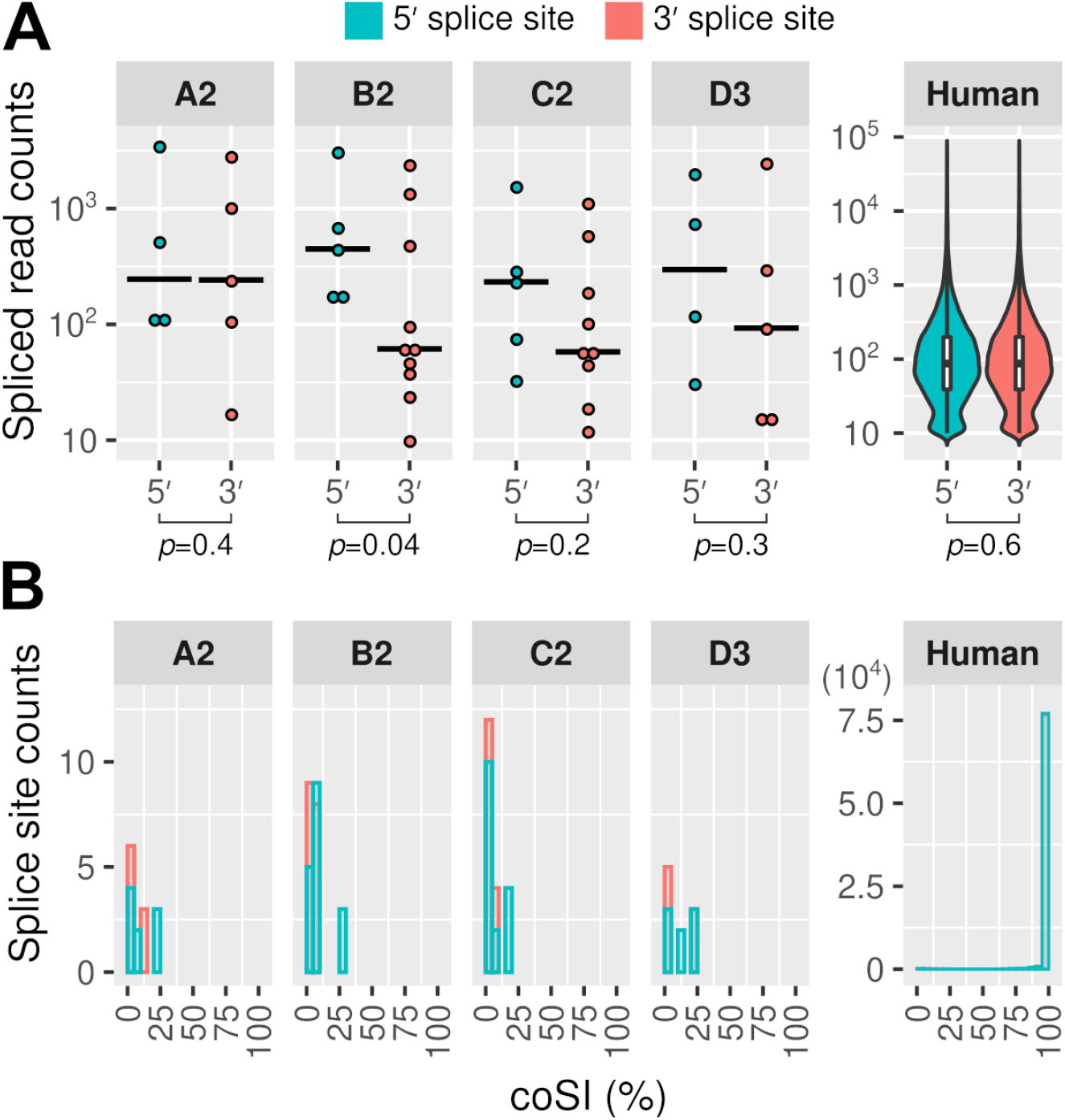
HBV 5′ splice sites are more likely to be spliced than 3′ splice sites. **(A)** More spliced reads were mapped across the 5′ splice sites of HBV than 3′ splice sites. Similar results were obtained from Welch two-sample t-test (one-sided) and permutation test (e.g. *p*-values of 0.04 and 0.06 were obtained for genotype B2, respectively). Solid black lines indicate median values. **(B)** Completeness of splicing at the 5′ and 3′ splice sites. Only the splice sites supported by at least 10 reads were included for comparison. coSI, completed Splicing Index.

To quantify the rates of splicing in HBV versus host cell RNAs, we calculated the completed Splicing Index (coSI) of the 5′ and 3′ splice sites. The 5′ splice sites of HBV showed higher coSI scores than 3′ splice sites (9% versus 5% on average; see also Fig 3B). In contrast, splicing was 87% and 86% completed at the host 5′ and 3′ splice sites, respectively. These results showed that the 5′ splice sites of HBV tend to be more frequently spliced than 3′ splice sites (e.g. see ② in Fig 2), but were much less efficiently utilised than host splice sites.

To identify the key differences between the HBV and human splice sites, we compared their splice site contexts using WebLogo (43), which used the frequencies of the uniquely mapped spliced reads to estimate the most frequently used splice sites. This approach showed that the nucleotide frequency distributions of human splice sites were similar to previous studies (Fig 4) (44). Notably, the most frequently used splice sites differed between the virus and host, e.g. −1 positions of the splice donor sites (Fig 4, left panel, U versus G shaded in gray). The differences between the HBV genotypes were marginal, as the spliced reads were predominantly mapped to SP1 and SP9 (Fig 1B and 2).

**Fig 4.**
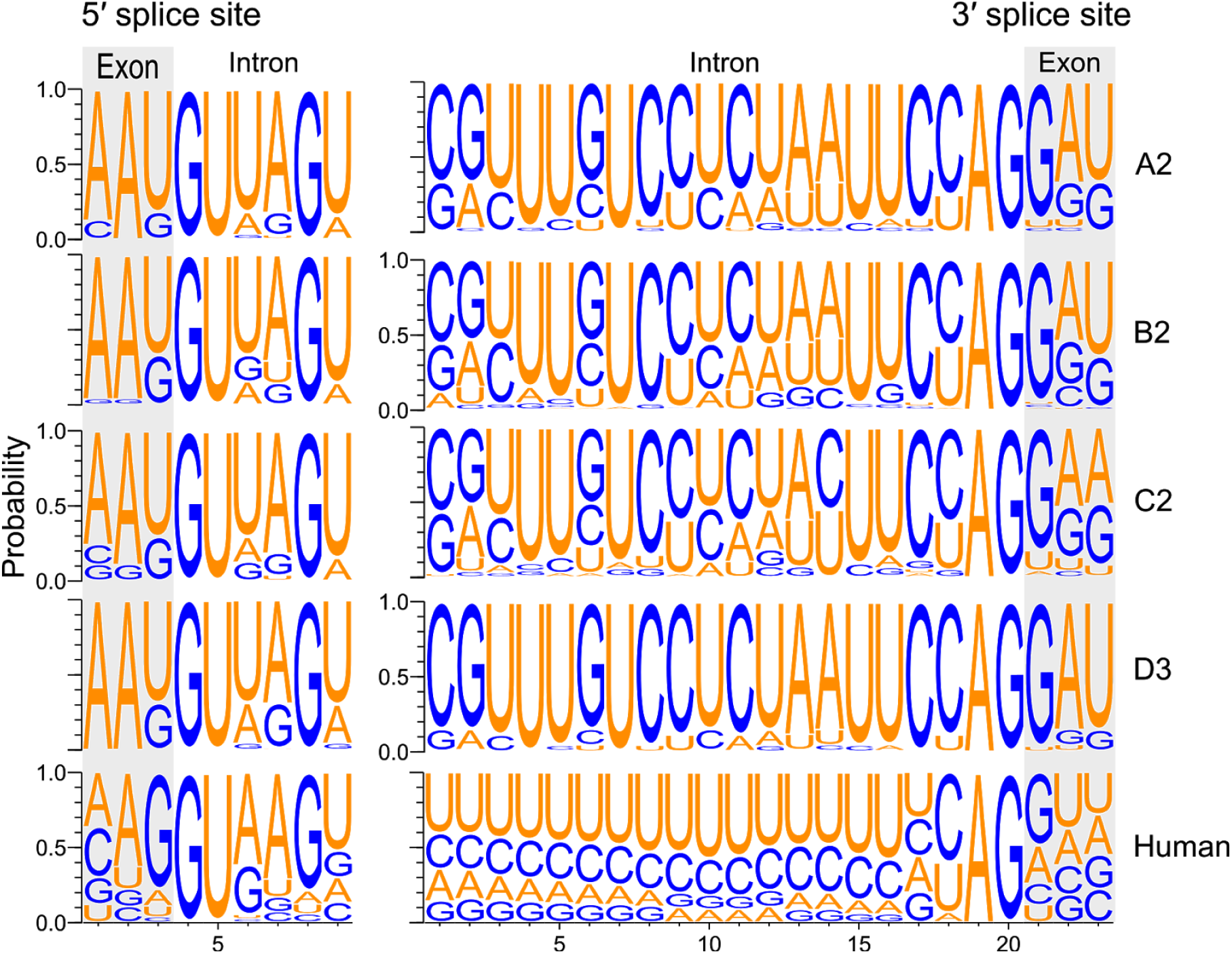
Most frequently used splice sites differed between HBV and the host. The nucleotide frequencies surrounding the splice sites are represented by the spliced reads. Exon boundaries are shaded in gray. Only the splice sites supported by at least 10 reads were included.

### HBV replication had little effect on the host gene expression

To understand the impact of HBV replication on the host, we carried out differential gene expression analysis using DESeq2 (45). We found that only one and 12 genes were significantly differentially expressed in A2 and B2 treated samples, respectively (Fig 5, red points; Fig 6 and Supplementary Table S6, FDR-adjusted *p*-value <0.05). Interestingly, both the A2 and B2 genotypes also showed higher levels of HBV transcriptomes (Fig S1; see also Supplementary Table S1 and S2). The accumulation of HBV transcripts may induce a stress response as the stress-related genes INHBE, FAM129A, SESN2, ASNS, and CHAC1 were all upregulated (Fig 7; see also the functional annotation results in Supplementary Table S7). Indeed, previous studies have also shown that HBV infection could lead to endoplasmic reticulum stress (46–48), including upregulation of INHBE (49). Interestingly, 3 significantly upregulated genes (ADM2, AKNA, and SH3BP2) were previously shown to correlate with the Ishak fibrosis stage (50).

**Fig 5.**
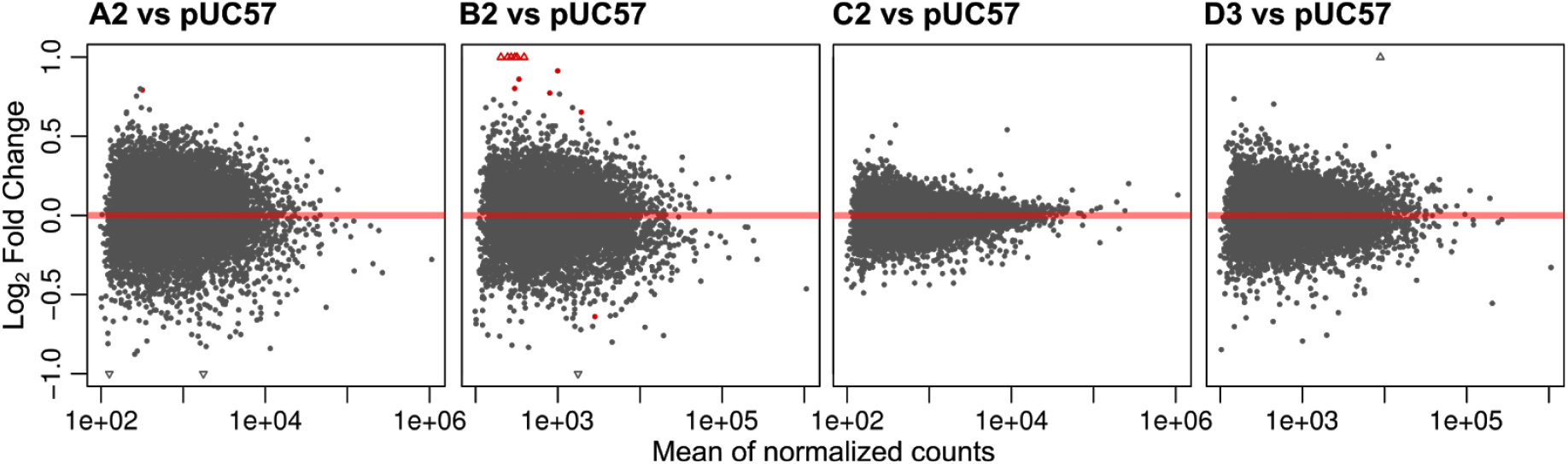
MA plots show differential gene expression between the HBV treated and control samples. Normalized counts indicate the counts divided by the normalization factors (as computed using the DESeq2 default function). Red points denote the FDR-adjusted *p*-value of <0.05. Unfilled triangles denote the genes that have undergone twofold changes in expression. MA plot, log ratio versus mean expression (log scale). See also the related Supplementary Table S6.

**Fig 6.**
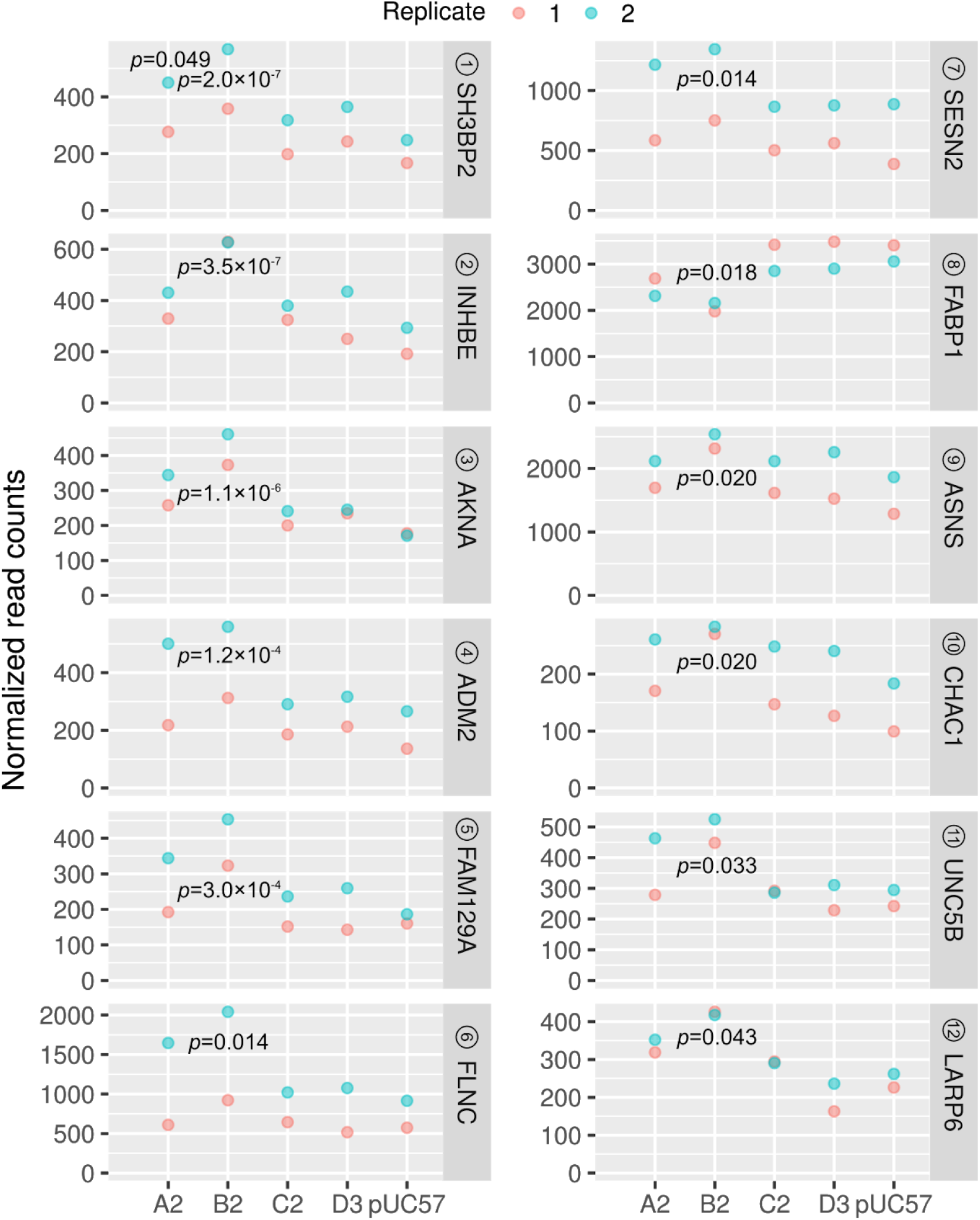
Significantly dysregulated genes in the HBV treated cells. A total of 12 genes were differentially expressed in the B2 treated sample. Circled numbers denote the ranking based on FDR-adjusted *p*-values. See also Supplementary Table S6 and S7.

**Fig 7.**
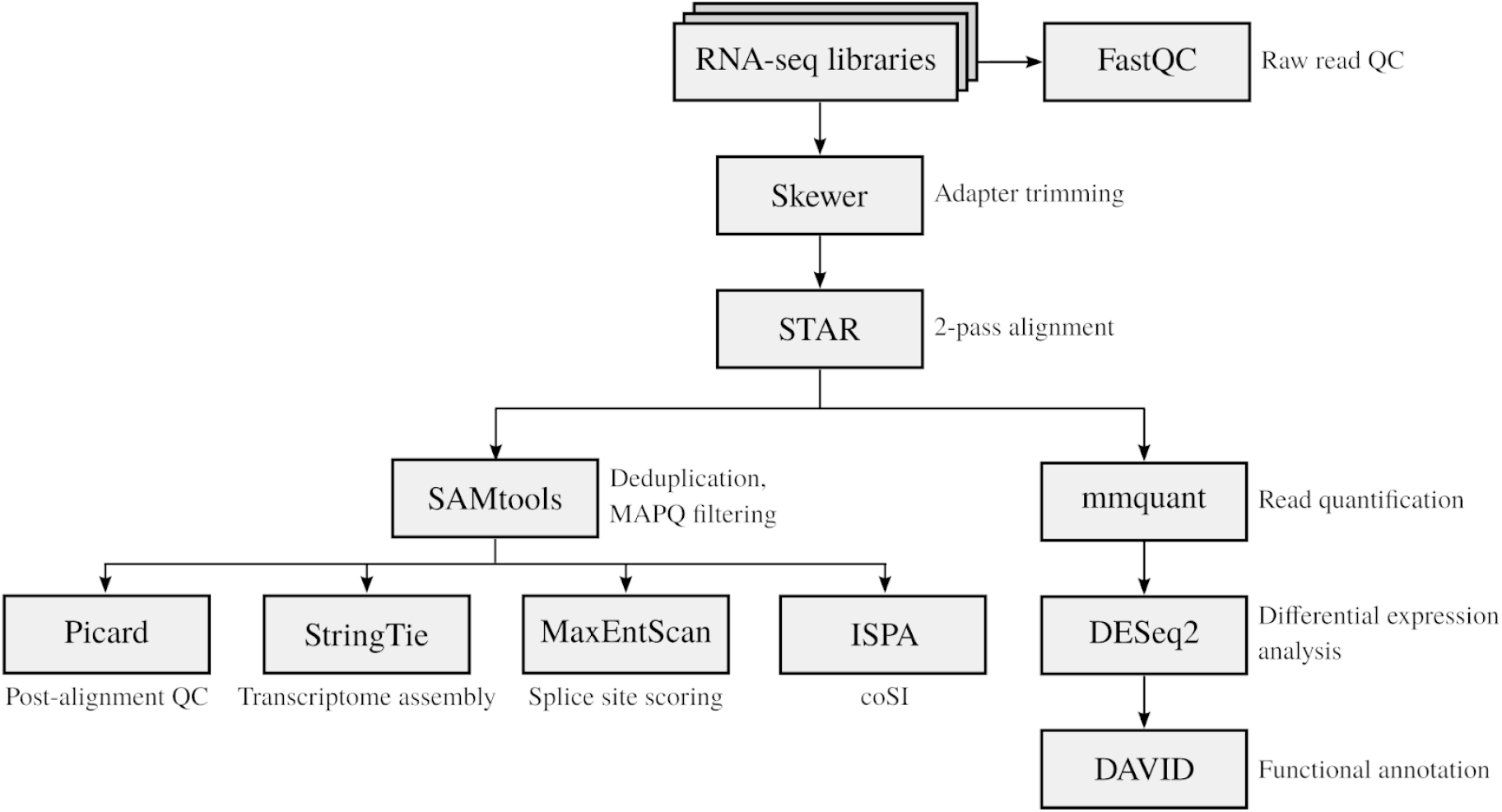
RNA-seq analysis of the HBV and host transcriptomes. Quality control (QC) check on the paired-end RNA-seq libraries was carried out using FastQC. Adapter sequences were trimmed from the RNA-seq reads using Skewer. Trimmed reads were aligned to the human and HBV genomes using STAR in 2-pass mode. Duplicated and multi-mapped reads were discarded from the Binary Alignment Map (BAM) files using SAMtools. Post-alignment QC check was performed using Picard tools. Transcriptome assembly and quantification were carried out using StringTie, with a post-processing step focusing on the HBV spliced transcript isoforms. Splice site sequence contexts were scored using MaxEntScan. Completeness of RNA splicing was evaluated using IPSA. Reads mapped to human genes were quantified using mmquant, followed by differential gene expression analysis using DESeq2. A list of differentially expressed genes was submitted to the DAVID webserver for functional annotation analysis. coSI, completed Splicing Index.

### Concluding remarks

Our study has used RNA-seq to identify the type and proportion of splice variants produced for HBV genotypes A2, B2, C2 and D3, with a higher degree of sensitivity than 454 sequencing and conventional reverse-transcription PCR (13, 14, 21, 24), all of which require an intervening step to generate cDNA, followed by PCR. We identified a number of novel splice variants across HBV genotypes as well as previously identified variants of yet unknown functions. The role of the SP9 variant in particular needs to be further explored. We acknowledge that this study is limited to one clone for each of the genotypes, and needs to be expanded to include additional HBV genotypes. Nonetheless, this study demonstrates that HBV has a large capacity for alternative splicing likely controlled by *cis*-acting elements such as the PRE (1), which results in high-levels of SP1 and SP9 mRNAs, despite the suboptimal context of the 5′ splice site. With up to 25% of all HBV mRNAs being of splice origin, the importance of these molecules in the HBV “life-cycle” and pathogenesis requires further investigation.

## MATERIALS AND METHODS

### Cell culture

Cell culture and transfection experiments were carried out as previously described with the following modifications (6). Huh7 cells were seeded in 6-well plates at partial confluence. After overnight incubation, the cells were transiently transfected with pUC57 constructs harboring 1.3-mer HBV genomes (genotypes A2, B2, C2, and D3) using FuGene 6 transfection reagent according to the manufacturer’s instructions (Promega, WI, USA). The generation of plasmids has been previously described and this transient expression system relies on the endogenous promoters of HBV for transcription (6). The empty pUC57 vector was used as a control. Two independent biological replicates were performed which included 2 technical replicates for each treatment.

### RNA-seq

Total RNA samples were purified using RNeasy kit (Qiagen, Hilden, Germany) and submitted to the Otago Genomics and Bioinformatics Facility at the University of Otago (Dunedin, New Zealand) under contract for library construction and sequencing. The libraries were prepared using TruSeq stranded Total RNA sample preparation kit with Ribo-Zero according to the manufacturer’s protocol and sequenced using HiSeq 2500 (Illumina, CA, USA), generating 125-bp paired-end reads (see the RNA-seq analysis workflow in Fig 7).

### Quality control (QC) of RNA-seq

The fastq files were examined using FastQC v0.11.5 (51). Most files passed most of the analysis modules except ‘per base sequence content’, ‘sequence duplication levels’, and ‘k-mer content’, which are common warnings for Illumina TruSeq reads (FastQC documentation). However, some fastq files failed at per base sequence quality and per base N content due to diminish of the quality score over position 100. Some fastq files also failed at per tile sequence quality due to loss of quality at random positions and cycles, which is likely due to the overloading of flowcell. Both of these issues should have minimum impact on downstream analysis because the regions of poor based calling were soft-clipped during alignment. In addition, only uniquely mapped reads were used for gene counting and transcript assembly.

As a post-alignment QC, the mapping statistics of the non-redundant RNA-seq reads were examined. About 60% of the reads were uniquely mapped reads to the human genomes (Supplementary Table S4). The distribution of aligned reads was then analyzed using the CollectRnaSeqMetrics mode of Picard 2.10.2 (http://broadinstitute.github.io/picard; Broad Institute, Cambridge, MA, USA). Over 55% and 27% of the bases of these reads were mapped the coding sequences (CDS) and untranslated regions (UTRs), respectively. Only 10% or lower of the bases of the sequencing reads were aligned to intronic or intergenic regions (Supplementary Table S5). These metrics are comparable with previous findings (52), indicating that our RNA-seq libraries are reliable.

### Sequence alignment

Adapter sequences were trimmed from the RNA-seq reads using Skewer v0.2.2 (53). To detect novel splice junctions, alignment was performed using STAR 2.5.2a in 2-pass mode (54). Duplicated reads were removed and uniquely mapped reads were retained using SAMtools (55).

### Transcriptome assembly

Transcriptome assembly of HBV splice variants was carried out using StringTie v1.3.3b (56). This tool assembles and quantifies spliced transcript isoforms using a network flow algorithm. Annotations of the spliced transcript isoforms were merged by biological replicates using gtfmerge (https://github.com/Kingsford-Group/rnaseqtools) and GffCompare (57). Only the assembled spliced transcripts that were found in both biological replicates were reported (intersection of complete, exact match intron chain). After merging the BAM files by biological replicates using SAMtools, a spliced graph of HBV transcripts was plotted using Gviz and GenomicFeatures (58, 59).

### Splice site analysis

Splice site sequence contexts were scored using MaxEntScan (60). This tool is a key plugin of the Ensembl Variant Effect Predictor (61) and performed the best in a recent benchmark (62). Splice site mapping frequencies were parsed from the SJ.out.tab file from STAR. IPSA was used to calculate the coSI score of 5′ and 3′ splice sites (https://github.com/pervouchine/ipsa) (63). WebLogo 3.5.0 was used to plot the nucleotide frequencies surrounding the splice sites.

### Differential gene expression analysis

To examine the reproducibility of the biological replicates, the uniquely mapped reads were first counted and summarized at the gene level using mmquant v1.3 (64). The correlation of samples was analyzed. The Spearman’s correlations between the biological replicates were >0.9, suggesting a good reproducibility (Supplementary Fig S2). However, the Spearman’s correlations between biological replicates were smaller than those within the same batch (e.g. A2_rep1 versus A2_rep2 is 0.938, whereas A2_rep1 versus B2_rep1 is 0.996). These results suggest the presence of batch effects, which is likely due to the second biological replicate being performed a year after. This was further examined using principal component analysis (PCA). Indeed, the samples were clustered by batches (Supplementary Fig S3).

To resolve the issue of batch effects, read counts were transformed using the vst (variance-stabilizing transformation) function of DESeq2 (45). Transformed read counts were examined using the plotPCA function of DESeq2, before and after correction using the removeBatchEffect function of limma (65). To take batch effects into account, differential expression analysis was carried out using batch as a linear term in the DESeqDataSetFromMatrix function. Differentially expressed genes were examined using DAVID functional annotation tools v6.8 (66, 67).

### Statistical analysis

Welch two-sample t-test and permutation test were performed using the exactRankTests R package (68, 69). Plotting was carried out using ggplot2 unless otherwise stated (70).

## Code and data availability

The raw RNA-seq libraries are available on Gene Expression Omnibus (71) (GSE155983). Scripts and data for the analysis can be found at https://github.com/lcscs12345/HBV_splicing_paper_2020

## ACKNOWLEDGMENTS

CMB and CSL were funded by the University of Otago. CSL is a recipient of a Dr. Sulaiman Daud 125th Jubilee Postgraduate Scholarship and the Marjorie McCallum travel award. PR and VS were funded by the NHMRC grant APP1145977.

## COMPETING INTERESTS

The authors declare that they have no competing interests.

